# Plant-based expression of SARS-CoV-2 antigen for use in an oral vaccine

**DOI:** 10.1101/2021.12.07.471131

**Authors:** Monique Power, Taha Azad, John C. Bell, Allyson M MacLean

## Abstract

Oral and intra-nasal vaccines represent a key means of inducing mucosal-based immunity against infection with SARS-CoV-2, yet such vaccines represent only a minority of candidates currently in development. In this brief communication, we assessed the expression of the SARS-CoV-2 Receptor Binding Domain (RBD) subunit of the surface-exposed Spike glycoprotein in the leaves of nine edible plant species (lettuce, spinach, collard greens, tomato, cucumber, radish, arugula, pepper, and Coho greens), with a goal of identifying a suitable candidate for the development of an oral vaccine against COVID-19. We report lettuce (*Lactuca sativa* L. cv. Hilde II Improved) to be a preferred host to support in planta expression of SARS-CoV-2 RBD, representing an important first step towards development of a plant-based oral vaccine.

## Main Text

While almost 300 vaccines against COVID-19 are presently in development, only a minority (3%) explore the potential of oral vaccines (WHO, 2021). Oral vaccines produced in plants offer several advantages over traditional processes, including potentially lower research and production costs, ease of scalability, and a negligible risk of contamination with human pathogens^1,2^, all of which are critical parameters towards facilitating equitable vaccine access to the world’s population.

By definition, a plant-based oral vaccine entails the expression of antigenic material within plants or green algae that, upon ingestion, elicits an immune response that is subsequently protective against disease^1^. A wide variety of plants and the model green alga *Chlamydomonas reinhardtii* have been explored for their potential to act as oral vaccines against an equally wide range of diseases. Examples involving human viruses include (and are not limited to) Hepatitis B in potato, tomato, lettuce, maize and *C. reinhardtii*^3–7^, HIV in tobacco, lettuce, carrot and *C. reinhardtii*^6,8–10^, HPV in tobacco, tomato and *C. reinhardtii*^11–13^, rabies virus in maize^14^, and influenza virus in maize^15^. Plant-based oral vaccines against bacterial and protozoal human pathogens have also been successfully created against, for example Enterotoxigenic *Escherichia coli* in rice^16^; *Mycobacterium tuberculosis* in carrot^17^; *Vibrio cholerae* in rice^18^, maize^19^ and potato^20^; *Plasmodium* in tobacco, lettuce ^21^, rapeseed^22^ and *C. reinhardtii*^23^; and *Toxoplasma gondii* in tobacco^24^. In an example that is highly relevant to our research, a subunit of the Spike protein of SARS-CoV (S1) was shown to induce SARS-CoV-specific IgA in mice after oral ingestion of transgenic tomato^25^.

The mechanisms by which plant-based oral vaccines elicit an immune response have been studied and reviewed in detail^1^. Ingested foods do not typically elicit an immune response, thus, to ensure efficient uptake by microfold cells within gut-associated lymphoid tissue, antigens may be fused to mucosal carrier proteins (e.g., B subunit of cholera toxin; CTxB). Plant-based oral vaccines have been shown to induce both cellular and humoral immunity within the mucosal and systemic immune systems^1,26^. Because the initial site of SARS-CoV-2 infection is within the respiratory mucosa, efforts to develop mucosal vaccines (oral or intra-nasal) are an important complement to conventional vaccines^2^.

This study aimed to optimize methods for the *in planta* expression of recombinant protein corresponding to the Receptor Binding Domain (RBD) subunit of the surface-exposed Spike glycoprotein of SARS-CoV-2. The RBD is a primary target of vaccine strategies against COVID-19, and most neutralizing antibodies against SARS-CoV-2 correspond to RBD^27,28^. We evaluated the suitability of nine edible plant species to express the viral antigen following transient transformation mediated by *Agrobacterium tumefaciens*, and then established parameters for optimal RBD expression in the most promising host plant.

Expression of recombinant proteins was trialed in lettuce (*Lactuca sativa* L. cv. Hilde II Improved), spinach (*Spinacia oleracea* L. cv. C2-608 Speedy Hybrid), collard greens (*Brassica oleracea* L. var. viridis), cucumber (*Cucumis sativus* L. cv Spartan Dawn F1 Hybrid), tomato (*Solanum lycopersicum* L. cv. Matina Organic), radish (*Raphanus sativus* L. cv. Cherry Belle), arugula (*Eruca vesicaria* L. (Cav.) ssp. *sativa* cv. Salad Rocket), peppers (*Capsicum annuum* L. cv. Purple Star Hybrid), and Choho Hybrid greens (*Brassica rapa* L. subsp. *narinosa* x *Brassica rapa* L. var. *perviridis*).

The plasmid vector selected for the expression of SARS-CoV-2 RBD *in planta* is pHREAC, which has been demonstrated to support a high level of protein expression in *Nicotiana benthamiana*^29^. The viral gene construct consists primarily of codon optimized (for expression in *N. benthamiana*) sequence corresponding to the RBD of the Spike glycoprotein (aa 329 to aa 522), fused to a cholera toxin B subunit (CTxB) to promote recombinant protein uptake across the intestinal epithelium. The expressed recombinant protein, RBD-CTxB, has a molecular mass of 40.3 kDa and includes HA- and FLAG-tags for antibody recognition (Fig.1a). Amongst the edible plants, the highest and most consistent expression of RBD-CTxB was found in lettuce (Fig.1b). Faint and/or inconsistent recombinant viral protein expression was observed in arugula, Choho Hybrid greens, pepper, spinach, and tomato. Arugula, Choho Hybrid greens and spinach showed no visible chlorosis or necrosis after infiltration, although tomato and pepper leaves developed both chlorosis and necrosis after infiltration. No expression of viral protein was observed in collard greens or radish, and leaves from these plants did not show visible damage from agroinfiltration. Lettuce was selected as the preferred host for edible vaccine production, as it consistently produced the highest expression of RBD-CTxB (based upon immunoblot analyses) and did not exhibit evidence of necrosis after agroinfiltration.

**Figure 1.**
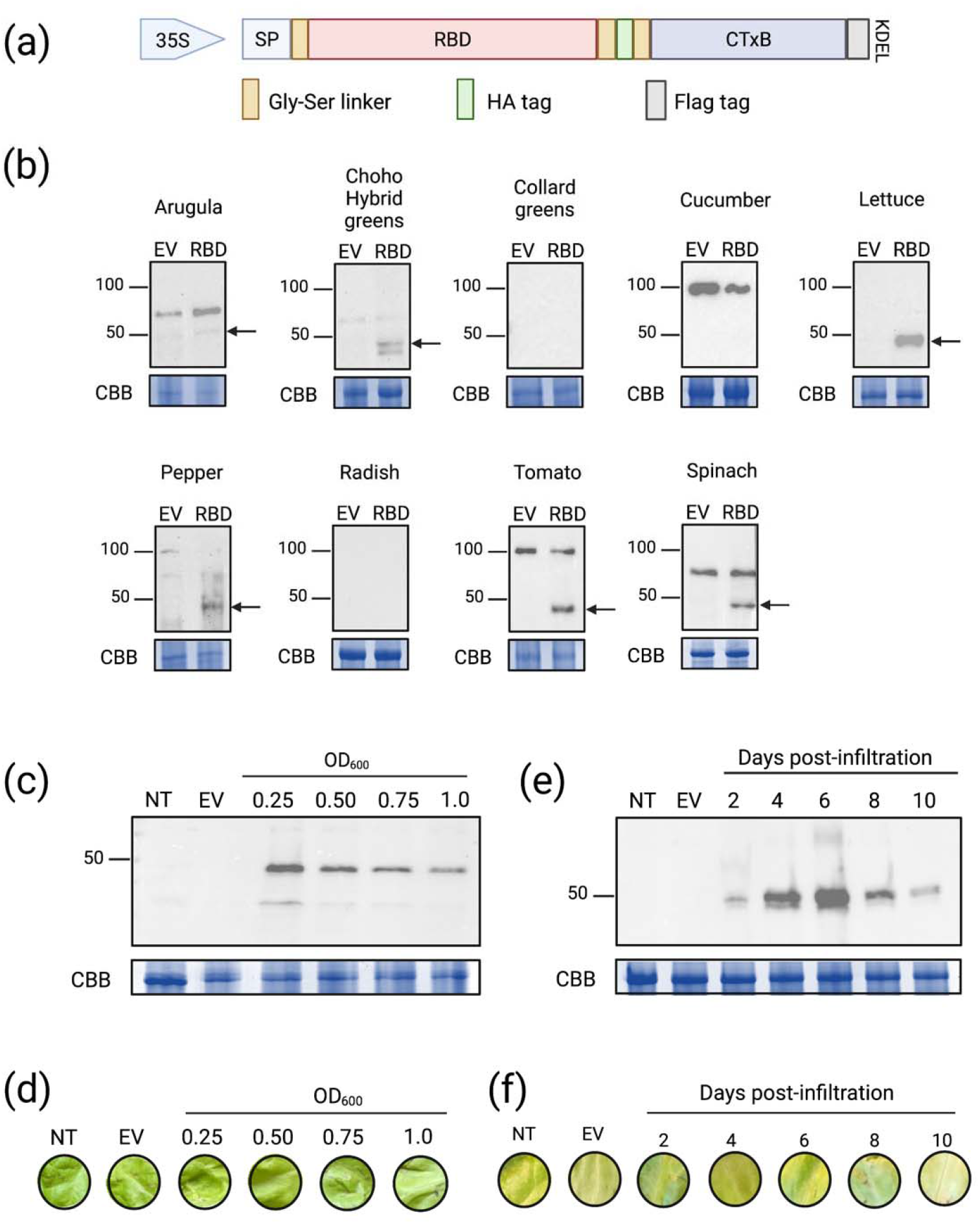
Optimization of expression of the Receptor Binding Domain (RBD) of SARS-CoV-2 Spike glycoprotein in an edible plant. (a) Schematic depiction of construct used to express RBD *in planta*. 35S, 35S CaMV promoter. SP, signal peptide PR-1a *Nicotiana tabacum* (aa 1 – 30). RBD corresponds to (aa 329 to aa 522) of the Wuhan-Hu-1 isolate of SARS-CoV-2. CTxB, B subunit of cholera toxin. KDEL, ER retention signal. (b) Anti-FLAG immunoblots to assess relative expression of RBD-CTxB recombinant protein in nine edible plant species. Arrows identify bands that correspond to the predicted molecular mass of RBD-CTxB (40.3 kDa). (c) Anti-FLAG immunoblot to assess RBD-CTxB expression following infiltration of lettuce with *A. tumefaciens* (RBD-CTxB/pHREAC), with adjusted OD600 as specified in panel. (d) Representative pictures of lettuce infiltrated with *A. tumefaciens* adjusted to OD600 as specified in panel. Leaf tissue was photographed 6 days post-infiltration of EV and RBD-CTxB. (e) Anti-FLAG immunoblot to assess RBD-CTxB expression over time, in lettuce infiltrated with an adjusted OD600 of 0.50. (f) Representative pictures of lettuce infiltrated with *A. tumefaciens* adjusted to an OD600 of 0.50. Leaf tissue was photographed 2 to 10 days post-infiltration of EV and RBD-CTxB, as indicated in panel. For panels (b to f), lettuce refers to *Lactuca sativa* L. cv. Hilde II Improved. NT, non-transformed lettuce collected (b) 4 days or (c to f) 6 days after infiltration of RBD-CTxB samples. EV, infiltration of *A. tumefaciens* (pHREAC), adjusted to OD600 of 0.50, collected (b) 4 days or (c to f) 6 days dpi. CBB, Coomassie Brilliant Blue stained SDS-PAGE to assess protein loading.

We next assessed the optimal parameters to maximize protein expression in this host. We observed comparable expression of RBD-CTxB between leaves infiltrated with an Agrobacterium suspension adjusted to an OD600 of 0.25 and 0.50, with minimal to no visible signs of chlorosis or necrosis (Fig.1c). Some yellowing was observed in tissue infiltrated at OD600 of 0.75 and 1.0, thus we decided to select an OD600 of 0.5 in subsequent experiments to minimize stress to the plant host (Fig.1d). We also monitored accumulation of RBD-CTxB at 2 to 10 days post-infiltration (dpi) to determine the timepoint at which expression is highest (Fig.1e) and damage to tissues was minimal (Fig.1f). RBD-CTxB expression appears to increase with time to achieve highest levels at 6 dpi, then declines from 8 dpi to 10 dpi. To obtain the highest possible expression of the protein of interest, our data indicate that lettuce plants should be harvested 6 days after infiltration.

In summary, this study represents one of the first protocols to express SARS-CoV-2 viral antigen in an edible plant species, a key step towards achieving the goal of developing a plant-based oral vaccine that is protective against COVID-19. Plant-based edible vaccines offer excellent potential towards enhancing equity in vaccine development and distribution in low-income developing countries, many of which report fewer than 2% of citizens as receiving even a single COVID-19 vaccine dose^30^.

## Athor Contributions

AMM and TA conceived and designed the project. MP performed all agroinfiltration experiments and immunoblots. MP and AMM contributed to the writing of the manuscript, with edits from TA and JCB. All authors reviewed and approved the manuscript.

## Acknowledgements

We are grateful for helpful discussions and advice from Rima Menassa and colleagues at Agriculture and Agri-Food Canada, Emmanuel Margolin, Edward Rybicki and colleagues from Biopharming Research Unit, University of Cape Town. We gratefully acknowledge the excellent support of Michelle Brazeau, University of Ottawa, in her management of our greenhouse. This study was supported in part by funding from the University of Ottawa Faculty of Science.

